# Inflammatory Monocytes Increase Prior to Detectable HIV-1 Rebound Viremia

**DOI:** 10.1101/2024.12.24.630257

**Authors:** Anna Farrell-Sherman, Natalia de la Force, Cecila Prator, Renan Valieris, Walker Azam, Israel Da Silva, Steven G. Deeks, Cassandra Thanh, Ronald Bosch, Timothy J. Henrich, Lillian Cohn

## Abstract

The persistence of HIV-1 proviruses in latently infected cells allows viremia to resume upon treatment cessation. To characterize the resulting immune response, we compare plasma proteomics and single-cell transcriptomics of peripheral blood mononuclear cells (PBMCs) before, during, and after detectable plasma viremia. We observe unique transcriptional signatures prior to viral rebound including a significant increase in CD16^++^ monocytes with increased anti-viral gene expression. Inflammatory proteins were identified in plasma after detectable rebound. Identifying early signals of imminent viral rebound after treatment cessation will aid in the development of strategies to prolong time to viral rebound and cure HIV-1.

## Introduction

Antiretroviral therapy (ART) prevents the infection of new cells by HIV-1 and prevents HIV-1 replication in people living with HIV (PLWH), however, viremia returns rapidly after treatment cessation. This process of viral rebound occurs due to a persistent reservoir of latently infected cells within which virus begins to replicate following treatment interruption [1]. We previously demonstrated that the frequency of CD4^+^ T cells expressing the Tumor Necrosis Factor Super Family Receptor, CD30, increases after ART is stopped but prior to detectable HIV-1 RNA in plasma [2]. Such a biomarker to predict HIV-1 viral rebound would greatly benefit HIV-1 cure studies.

We hypothesize that the immune system responds to low-level viral activity before it is detectable by clinical assays, potentially generating signals that could serve as an indicator for imminent rebound. Here, we characterize immune signatures during treatment interruption of 24 PLWH who participated in the placebo arm of three separate ATCG vaccine trials [3-5]. We performed high-dimensional plasma proteomics on samples from 23 participants and single-cell RNAseq on 10 participants who were chosen because they demonstrated the greatest fold change in CD4^+^ T cell CD30 expression after stopping ART in our prior study. Examining the viral-immune dynamics during rebound is essential to understanding mechanisms of HIV immune control.

## Methods

Cohort: We assembled a cohort of 24 individuals who participated in observational analytical treatment interruption (ATI) studies as part of three previously published ACTG trials (ACTG5068, ACTG5024, ACTG5187) [3-5]. Two of the three trials enrolled participants who were treated in the acute or early phase of HIV-1, and one enrolled ART naïve participants who were treated for at least 44 weeks before undergoing ATI. Consistent with the parent ACTG trials, 21 (87.5%) of participants were male and the median age was 42 years (IQR 37-48). Fourteen participants were on an NNRTI-based regimen, 6 were on a protease inhibitor-based regimen, 3 were taking both PI and NNRTIs together and 1 was on NRTI only regimen.

Plasma Proteomics: We performed high-dimensional plasma proteomic analysis through three Olink targeted panels: inflammation, immuno-oncology, and biological processes with samples from 23 of our 24 participants. The median time to detected rebound (>50 copies/mL) in this cohort was 34 days (range 19-63). The median time from the initiation of ATI (the time of stopping ART) to pre-rebound sampling was 25 days (range 11-45).

These panels analyzed the abundance of 188 soluble proteins (Supplementary Table 1) in peripheral plasma at each of our three study timepoints. Differences in plasma protein abundance were assessed with a paired Wilcoxon test and p-values were corrected for False Discovery Rates (FDR).

Single-cell RNA sequencing (scRNAseq): A subset of 10 participants were selected based on previously published high fold changes in CD30 surface expression on CD4^+^ T cells between samples collected on-ART and pre-rebound. The median time to rebound in this cohort was 33 days (range 19-51). The median time from the initiation of ATI (the time of stopping ART) to pre-rebound sampling was 25 days (range 13-43). Live peripheral blood mononuclear cells (PBMCs) were processed through the 10x Genomics 5’ Single Cell assay on a 10x Genomics chip K according to manufacturer’s instructions. Library products were sequenced on the Illumina NovaSeq platform with a depth of 17068-92846 reads per cell after filtering. Three libraries failed, resulting in a total of 27 samples from 10 donors across 3 time points. Libraries were processed with Cell Ranger 6.1.1. Cells were removed if they were associated with less than 2500 transcripts, fewer than 900 genes, or if mitochondrial gene expression comprised more than 15% of the reads.

scRNAseq Analysis: Single cell data was analyzed with Seurat v4 in R. Reads in each cell were normalized by cell cycle phase and mitochondrial expression ratio, then cell type was estimated by mapping to a standard PBMC multi-modal dataset [6]. We selected the top 3000 most variable genes for integration, with on-ART samples as reference, to determine differentially expressed genes. Cell-type abundance was tested with a non-parametric Wilcoxon signed-rank test paired by participant to account for baseline differences (with FDR corrected p-values). Cell expression counts were summed by sample and by cell type to form pseudo-bulk count matrices which were analyzed with DESeq2 [7]. The results of the differential expression analysis were used to rank the genes and create a subset of 219 differentially expressed genes (unadjusted p-value < 0.05). Finally, pathways enriched in this subset were identified with gene set enrichment analysis, implemented by the clusterProfiler R package [8]. We evaluated hallmark, curated and gene ontology gene sets from MSigDB [9].

## Results

To investigate the immune-viral interactions of rebound, we compared samples from three distinct time points: on-ART (before ATI), pre-rebound (during ATI, before detectable plasma viremia), and post-rebound (during ATI, detectable viremia) (Figure 1A). Cryopreserved PBMCs and plasma samples were obtained at these three time-points from each participant. Proteomic analyses revealed a significant increase in plasma LAG3, TNF, GZMH, CRTAM, CXCL10, IL12, TRAIL, CD27, MIC-A/B, CD5, MMP12, GZMB, IL-18R1, CXCL9, PD-L2, GZMA, NCR1, CD83, SLAMF1, AND IL-12B from on-ART to post-viral rebound (FDR adjusted p-values <0.05; Figure 1B, Table S1), but no significant differences were observed between on-ART and pre-rebound time points (FDR >0.05).

**FIGURE 1:**
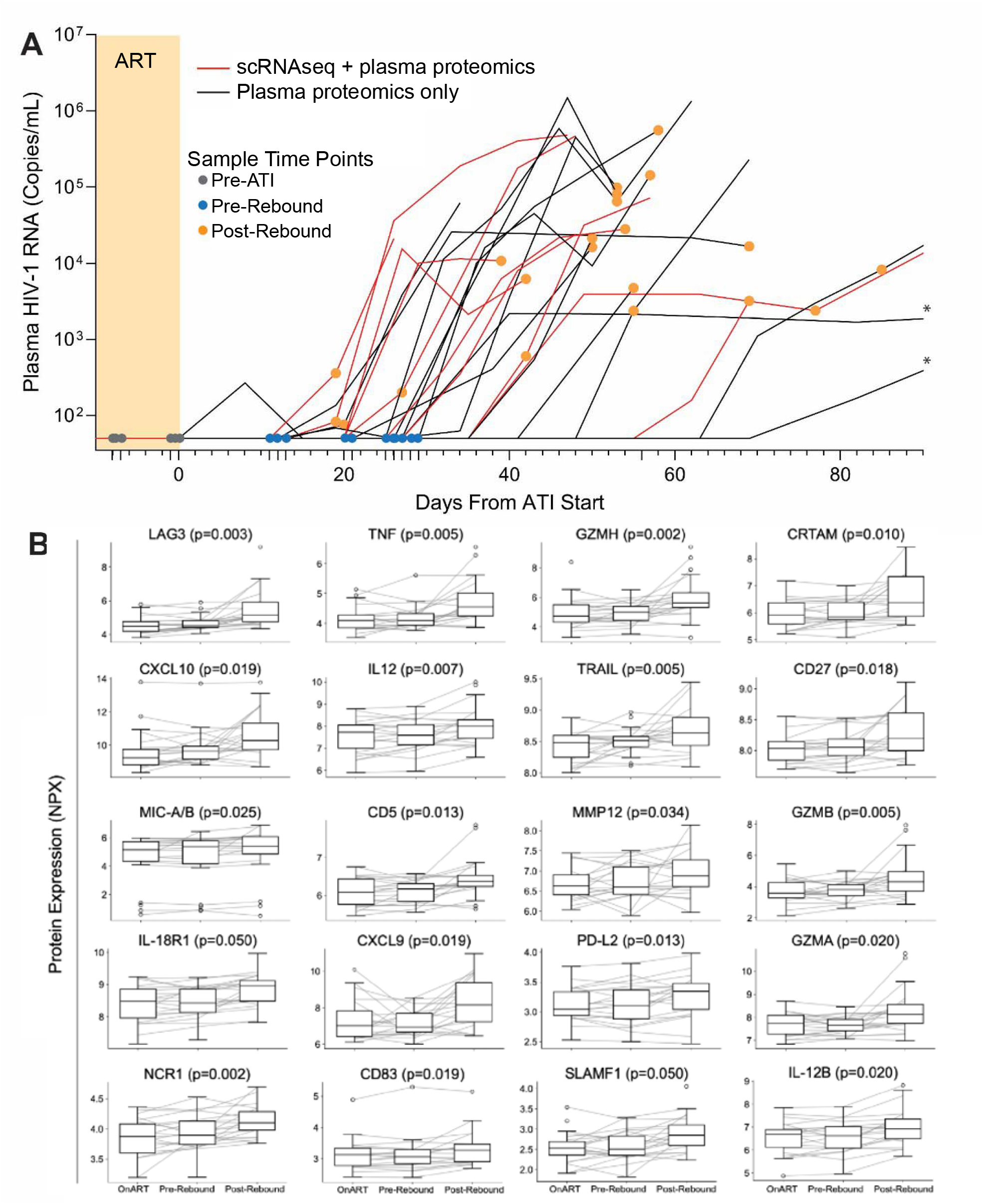
(A) Plasma viral load levels from all 24 participants included in the proteomics and scRNAseq experiments throughout ATI. Colored circles indicate samples used in the current study. Yellow shading indicates ART. The viral load limit of detection is 50 HIV-1 RNA copies/mL. Lines with asterisks indicate post-rebound sampling >90 days from first detectable viral load measurements. (B) Circulating proteins significantly upregulated between on-ART and post-rebound timepoints as detected by O-link (full results are show in Table S1). P-values are from paired comparisons of on-ART to post-rebound protein levels.

Next, we performed 10x scRNAseq on 10 participants (8 participants had both an on-ART and pre-rebound time point) (Figure 2) and analyzed cell type proportions. The median proportion of classical CD14^++^ monocytes increased from 16.6% of total PBMCs to 19.0% (p=0.054) and we observed a significant increase in the median proportion of anti-viral CD16^++^ monocytes from on-ART baseline to pre-rebound (2.57% to 4.38% of PBMC; p=0.008) (Figure 2AB). The baseline on-ART levels of CD16^++^ monocytes varied from 1-5%, across donors, however, the increase in CD16^++^ monocytes remained consistently ∼1.7x. No other subsets significantly differed in abundance from on-ART to pre-rebound.

**FIGURE 2:**
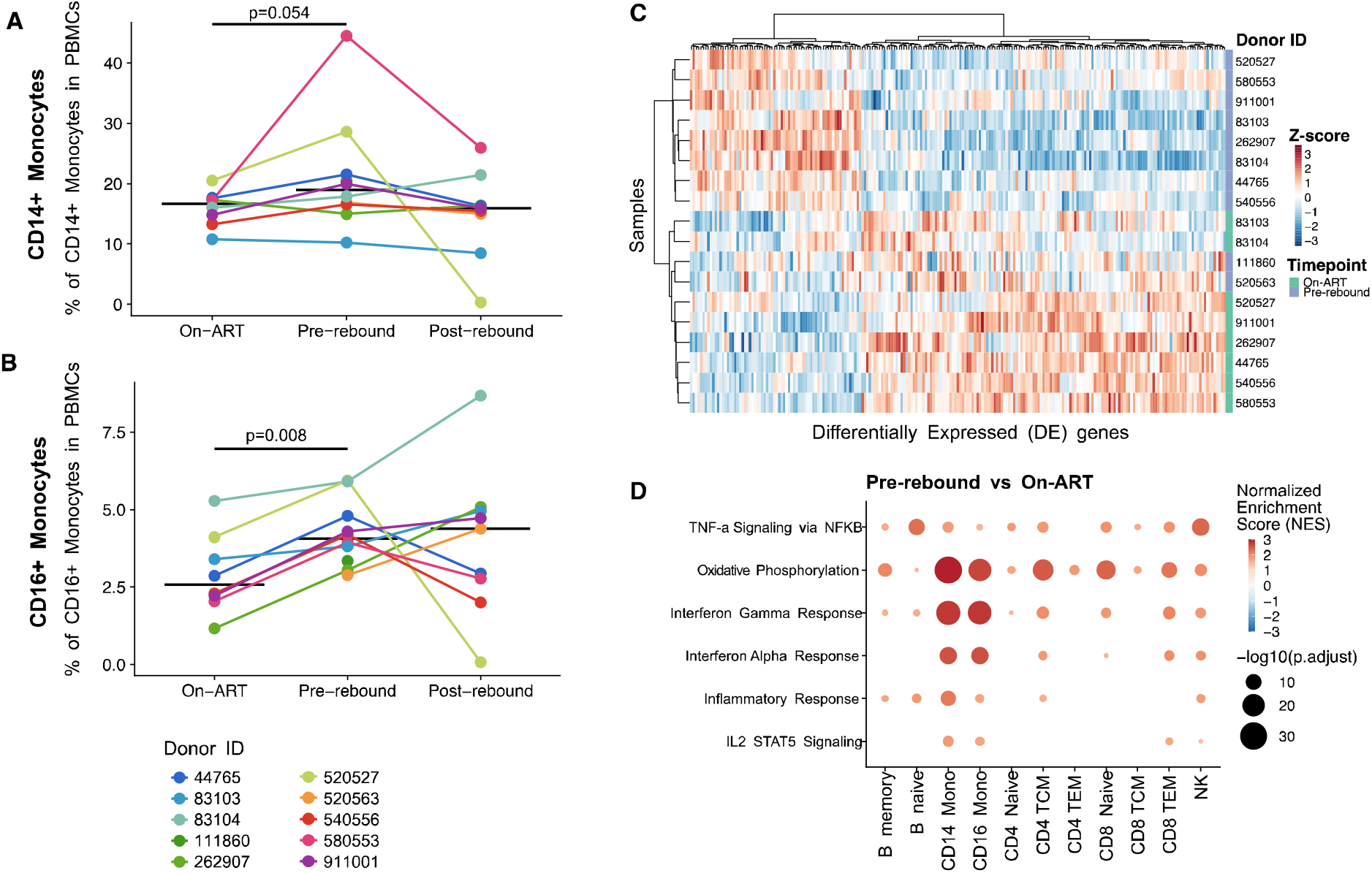
(A-B) Changes in CD14^++^ (A) and CD16^++^ (B) cell type frequencies across timepoints. Individual donors shown in color with the median indicated with a horizontal black line. P-values from a Wilcoxon test between indicated timepoints. (C) Heatmap of the z-score of normalized expression for significantly differentially expressed genes in CD16^++^ monocytes between on-ART and pre-rebound (significance before multiple hypothesis testing). (D) Changes in pathway expression between on-ART and pre-rebound in various cell types. Bubbles are colored by normalized enrichment score (NES) and the size indicates the significance of the association -log10(p-value).

To assess changes in gene expression between cells on-ART and pre-rebound, we performed a pseudo-bulk differential gene expression analysis on our scRNAseq data. In CD16^++^ monocytes, there were 219 differentially expressed genes which clearly distinguish the on-ART and pre-rebound expression pattern of these cells (Figure 2C). Pathway analysis on the differentially expressed genes from monocyte subsets and other cell type subsets show upregulation of specific inflammatory pathways, including TNF_α_ via NF_K_B, oxidative phosphorylation, interferon response, inflammation, and IL2/STAT5 signaling (Figure 2D). Whereas expression of these pathways increased in multiple cell subsets, we observed the greatest fold change differences in monocyte subsets.

## Discussion

In this study, we evaluate changes in the immune landscape of PLWH throughout ATI to determine whether an immune response is detectable in peripheral blood prior to clinical viral rebound. While we only identify increases in inflammatory circulating proteins following HIV-1 recrudescence, we observe a significant increase in CD16^++^ monocytes and a significant increase in the expression of inflammatory pathways in monocyte subsets before HIV-1 RNA was detected in plasma.

Monocytes are comprised of three subsets: classical, intermediate, and non-classical, however intermediate monocytes cannot be differentiated with current single-cell RNA sequencing analyses [10]. Classical monocytes, which make up ∼80% of the monocyte compartment, express the LPS receptor CD14 but not CD16, while both intermediate and non-classical subsets express CD16 [11]. Intermediate monocytes maintain CD14 expression, while non-classical monocytes are CD14^dim^CD16^++^ and functionally distinct from the other subsets with higher expression of anti-viral genes and a more migratory phenotype, which helps alert the immune system to viral replication in tissues [11, 12]. Migration throughout the body after detecting HIV-1 activity in tissues could explain why this subset is measurable before detection of viremia in our study.

Strikingly, the increase in anti-viral CD16^++^ monocyte frequency was consistent across all participants, at a median of 13 days prior to the first detectable viral load measurement. Previous studies suggest that non-classical monocytes are a first-line immune response to viruses including HIV-1 which increase in frequency early after HIV acquisition and wax and wane in concordance with viral load through all stages of HIV progression [12-14]. These data suggest that monocytes may be a relatively universal responsive to HIV-1 activation in localized tissues before systemic viral replication, with non-classical CD16^++^ monocytes being particularly sensitive to viral activity.

The increasing expression of specific inflammatory pathways within expanding monocyte subsets supports the hypothesis that these cells not only increase in frequency but may be engaged in suppressing viral replication in tissues. In fact, both classical CD14^++^ and CD16^++^ monocytes were highly enriched in inflammatory pathways and responses related to IFN-y and IFN-_α_. These pathways are an integral part of the antiviral response and are necessary for the production of cytokines to recruit immune cells to sites of viral replication [15]. In addition, CD14^++^ and CD16^++^ monocytes increase expression of pathways related to oxidative phosphorylation, indicating that energy requirements may increase in response to proliferation and pathogenesis. Future studies more deeply investigating inflammatory anti-viral monocyte subsets and further tissue-based studies during ATI are urgently needed.

Our study has a number of potential implications for HIV cure, most importantly in identifying a signal of imminent viral rebound. Although sustained post-ART control is now being observed, there is no simple way to monitor these viral controllers for eventual viral rebound [16]. If current efforts to achieve durable virus control succeed, then monitoring individuals’ rebound kinetics will become paramount [17]. An assay that detects early virus spread would be particularly useful, as we suggested in an earlier case study of sustained post-ART control [18]. Our data also have implications for the development of novel interventions. Theoretically, the early immune response to replicating HIV following treatment interruption might prove to be an essential factor in determining who controls versus who does not control their virus [19].

Our study also demonstrates the need for more in-depth ATI studies examining immune responses during treatment interruption in response to rebound viremia. While the current study is based on a small number of participants sampled relatively infrequently during ATI, our results suggest that future studies with more frequent longitudinal sampling, and, if possible, intensive tissue sampling would greatly benefit the field. These studies would have the potential to examine early viral activity in tissues, to explain variation in time to HIV-1 rebound, and shed light into potential etiologies of post-treatment and post-intervention control of viremia.

## Acknowledgments

We thank the participants, staff, and principal investigators of the AIDS Clinical Trial Group (ACTG) studies A371 (Paul Volberding and Elizabeth Connick), A5024 (J. Michael Kilby and Ronald Mitsuyasu), A5068 (Jeffrey Jacobson, Ian Frank, Michael Saag, and Joseph Eron), A5187 (Daniel Barouch, Eric Rosenberg, and Daniel Kuritzkes), and A5197 (Robert Schooley, Michael Lederman, and Diane Havlir).

## Disclaimer

The content is solely the responsibility of the authors and does not necessarily represent the official views of the National Institutes of Health.

## Financial Support

This research was funded by the National Institute of Allergy and Infectious Diseases (Grants K24AI174971 to TJH, R01AI141003 to TJH, UM1AI164560 to SGD), and by the Bill and Melinda Gates Foundation (INV-002707 to LBC). This research was additionally supported by the Genomics & Bioinformatics Shared Resource, RRID:SCR_022606, of the Fred Hutch/University of Washington/Seattle Children’s Cancer Consortium (P30 CA015704).

## Potential conflicts of interest

T. J. H. received grant support from Merck and has consulted for Roche outside of this work.

