## Supplemental Material for "Inflammatory Monocytes Increase Prior to Detectable HIV-1 Rebound Viremia"

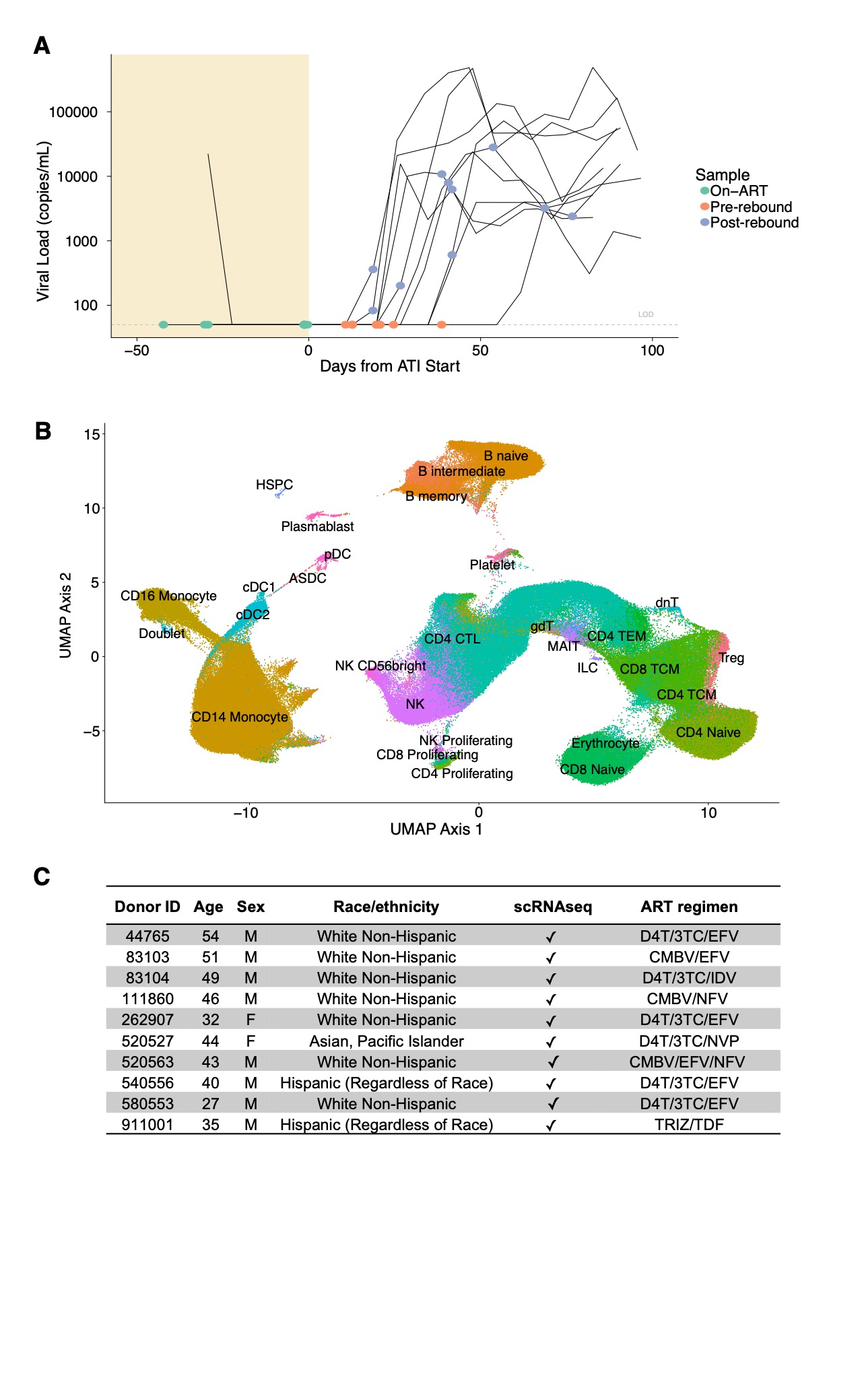

**Supplementary Figure 1.**  scRNAseq visualized with a UMAP projection based on the normalized RNA expression of the top 3000 variable genes. scRNAseq data was generated in a subset of 10 participants prior to and during ATI (prior to viral rebound) and following HIV-1 RNA detection in the plasma (viral recrudescence).

**Supplementary Table 1.** Soluble proteins measured by the Olink proteomic platform in plasma from participants at each sample time point. P-values from a Wilcoxon signed-rant test between on-ART and post-rebound protein concentration (NPX).

|  | **GeneID** | | | **On-ART vs Post-Rebound Wilcoxon Test** | | |
| --- | --- | --- | --- | --- | --- | --- |
| **Panel** | **Gene Name** | **OlinkID** | **Uniprot ID** | **Test Statistic** | **p-value** | **FDR Corrected p-value** |
| OT_96I | TNF | OID05548 | P01375 | 20 | 0.0004 | 0.0196 |
|  | IL-12B | OID00523 | P29460 | 21 | 0.0004 | 0.0196 |
|  | TRAIL | OID00488 | P50591 | 30 | 0.0019 | 0.0503 |
|  | SLAMF1 | OID00502 | Q13291 | 32 | 0.0025 | 0.0503 |
|  | IL-18R1 | OID00517 | Q13478 | 34 | 0.0033 | 0.0503 |
|  | CXCL10 | OID00535 | P02778 | 34 | 0.0033 | 0.0503 |
|  | CXCL9 | OID00490 | Q07325 | 38 | 0.0055 | 0.0594 |
|  | CD5 | OID00531 | P06127 | 39 | 0.0063 | 0.0594 |
|  | TNFRSF9 | OID00553 | Q07011 | 39 | 0.0063 | 0.0594 |
|  | IL-15RA | OID00514 | Q13261 | 40 | 0.0071 | 0.0594 |
|  | IL10 | OID00528 | P22301 | 40 | 0.0071 | 0.0594 |
|  | EN-RAGE | OID00541 | P80511 | 47 | 0.0158 | 0.1210 |
|  | CD8A | OID05124 | P01732 | 48 | 0.0175 | 0.1242 |
|  | uPA | OID00481 | P00749 | 51 | 0.0239 | 0.1568 |
|  | CCL19 | OID00513 | Q99731 | 53 | 0.0290 | 0.1780 |
|  | IL6 | OID00482 | P05231 | 55 | 0.0351 | 0.2016 |
|  | TRANCE | OID00521 | O14788 | 56 | 0.0384 | 0.2080 |
|  | TWEAK | OID00555 | O43508 | 57 | 0.0421 | 0.2151 |
|  | CSF-1 | OID00562 | P09603 | 60 | 0.0547 | 0.2586 |
|  | MCP-1 | OID00484 | P13500 | 61 | 0.0595 | 0.2586 |
|  | CDCP1 | OID00476 | Q9H5V8 | 62 | 0.0646 | 0.2586 |
|  | CD6 | OID00499 | P30203 | 62 | 0.0646 | 0.2586 |
|  | DNER | OID01213 | Q8NFT8 | 62 | 0.0646 | 0.2586 |
|  | IFN-gamma | OID05547 | P01579 | 64 | 0.0760 | 0.2689 |
|  | NT-3 | OID00554 | P20783 | 64 | 0.0760 | 0.2689 |
|  | TNFB | OID00561 | P01374 | 64 | 0.0760 | 0.2689 |
|  | TGF-alpha | OID00503 | P01135 | 65 | 0.0822 | 0.2701 |
|  | PD-L1 | OID00518 | Q9NZQ7 | 65 | 0.0822 | 0.2701 |
|  | IL-2RB | OID00492 | P14784 | 66 | 0.0888 | 0.2817 |
|  | CX3CL1 | OID00552 | P78423 | 67 | 0.0958 | 0.2877 |
|  | MCP-4 | OID00504 | Q99616 | 68 | 0.1032 | 0.2877 |
|  | MMP-1 | OID00510 | P03956 | 68 | 0.1032 | 0.2877 |
|  | MCP-2 | OID00549 | P80075 | 68 | 0.1032 | 0.2877 |
|  | ST1A1 | OID00557 | P50225 | 69 | 0.1111 | 0.3005 |
|  | STAMBP | OID00558 | O95630 | 70 | 0.1193 | 0.3081 |
|  | GDNF | OID00475 | P39905 | 71 | 0.1281 | 0.3081 |
|  | SIRT2 | OID00538 | Q8IXJ6 | 71 | 0.1281 | 0.3081 |
|  | CCL25 | OID00551 | O15444 | 71 | 0.1281 | 0.3081 |
|  | FGF-5 | OID00509 | P12034 | 72 | 0.1373 | 0.3081 |
|  | IL-10RB | OID00515 | Q08334 | 72 | 0.1373 | 0.3081 |
|  | 4E-BP1 | OID00536 | Q13541 | 72 | 0.1373 | 0.3081 |
|  | IL-24 | OID00524 | Q13007 | 58 | 0.1447 | 0.3144 |
|  | CCL23 | OID00530 | P55773 | 73 | 0.1470 | 0.3144 |
|  | CXCL11 | OID00486 | O14625 | 74 | 0.1571 | 0.3213 |
|  | IL4 | OID00546 | P05112 | 74 | 0.1571 | 0.3213 |
|  | AXIN1 | OID00487 | O15169 | 75 | 0.1678 | 0.3356 |
|  | IL-20RA | OID00489 | Q9UHF4 | 77 | 0.1907 | 0.3733 |
|  | IL7 | OID00478 | P13232 | 78 | 0.2029 | 0.3810 |
|  | CCL11 | OID00505 | P51671 | 78 | 0.2029 | 0.3810 |
|  | ARTN | OID00526 | Q5T4W7 | 79 | 0.2157 | 0.3969 |
|  | OPG | OID00479 | O00300 | 80 | 0.2290 | 0.3975 |
|  | IL-1 alpha | OID00493 | P01583 | 80 | 0.2290 | 0.3975 |
|  | IL-22 RA1 | OID00516 | Q8N6P7 | 80 | 0.2290 | 0.3975 |
|  | FGF-23 | OID00507 | Q9GZV9 | 82 | 0.2572 | 0.4226 |
|  | Flt3L | OID00533 | P49771 | 82 | 0.2572 | 0.4226 |
|  | CASP-8 | OID00550 | Q14790 | 82 | 0.2572 | 0.4226 |
|  | LIF | OID00547 | P15018 | 83 | 0.2722 | 0.4394 |
|  | CXCL5 | OID00520 | P42830 | 84 | 0.2877 | 0.4564 |
|  | CXCL6 | OID00534 | P80162 | 87 | 0.3377 | 0.5265 |
|  | IL-17C | OID00483 | Q9P0M4 | 88 | 0.3554 | 0.5450 |
|  | CD244 | OID00477 | Q9BZW8 | 91 | 0.4120 | 0.6213 |
|  | CCL28 | OID00539 | Q9NRJ3 | 92 | 0.4319 | 0.6409 |
|  | CXCL1 | OID00496 | P09341 | 94 | 0.4733 | 0.6804 |
|  | IL33 | OID00543 | O95760 | 94 | 0.4733 | 0.6804 |
|  | IL-17A | OID00485 | Q16552 | 95 | 0.4948 | 0.6897 |
|  | IL5 | OID00559 | P05113 | 95 | 0.4948 | 0.6897 |
|  | MCP-3 | OID00474 | P80098 | 96 | 0.5168 | 0.6991 |
|  | CCL20 | OID00556 | P78556 | 96 | 0.5168 | 0.6991 |
|  | LAP TGF-beta-1 | OID00480 | P01137 | 97 | 0.5392 | 0.7086 |
|  | FGF-21 | OID00512 | Q9NSA1 | 97 | 0.5392 | 0.7086 |
|  | TSLP | OID00497 | Q969D9 | 99 | 0.5854 | 0.7480 |
|  | TNFSF14 | OID00506 | O43557 | 99 | 0.5854 | 0.7480 |
|  | NRTN | OID00548 | Q99748 | 100 | 0.6091 | 0.7677 |
|  | CCL3 | OID00532 | P10147 | 101 | 0.6333 | 0.7754 |
|  | IL8 | OID00471 | P10145 | 102 | 0.6578 | 0.7754 |
|  | IL2 | OID00495 | P60568 | 102 | 0.6578 | 0.7754 |
|  | CCL4 | OID00498 | P13236 | 102 | 0.6578 | 0.7754 |
|  | Beta-NGF | OID00519 | P01138 | 102 | 0.6578 | 0.7754 |
|  | VEGFA | OID00472 | P15692 | 103 | 0.6827 | 0.7754 |
|  | OSM | OID00494 | P13725 | 104 | 0.7079 | 0.7754 |
|  | IL-10RA | OID00508 | Q13651 | 104 | 0.7079 | 0.7754 |
|  | IL13 | OID00525 | P35225 | 104 | 0.7079 | 0.7754 |
|  | IL-20 | OID00537 | Q9NYY1 | 104 | 0.7079 | 0.7754 |
|  | CD40 | OID00542 | P25942 | 104 | 0.7079 | 0.7754 |
|  | CST5 | OID00491 | P28325 | 107 | 0.7854 | 0.8402 |
|  | FGF-19 | OID00545 | O95750 | 107 | 0.7854 | 0.8402 |
|  | HGF | OID00522 | P14210 | 108 | 0.8117 | 0.8583 |
|  | SCF | OID00500 | P21583 | 110 | 0.8649 | 0.9042 |
|  | LIF-R | OID00511 | P42702 | 112 | 0.9187 | 0.9391 |
|  | ADA | OID00560 | P00813 | 112 | 0.9187 | 0.9391 |
|  | MMP-10 | OID00527 | P09238 | 113 | 0.9457 | 0.9561 |
|  | IL18 | OID00501 | Q14116 | 115 | 1.0000 | 1.0000 |
| OT_96IO | GZMH | OID00783 | P20718 | 7 | 0.0000 | 0.0017 |
|  | NCR1 | OID00816 | O76036 | 10 | 0.0000 | 0.0019 |
|  | LAG3 | OID05553 | P18627 | 14 | 0.0001 | 0.0032 |
|  | TRAIL | OID00769 | P50591 | 18 | 0.0002 | 0.0054 |
|  | TNF | OID05554 | P01375 | 20 | 0.0004 | 0.0054 |
|  | GZMB | OID00840 | P10144 | 20 | 0.0004 | 0.0054 |
|  | IL12 | OID00842 | P29459.P29460 | 22 | 0.0005 | 0.0067 |
|  | CRTAM | OID00766 | O95727 | 25 | 0.0009 | 0.0098 |
|  | CD5 | OID00812 | P06127 | 28 | 0.0014 | 0.0126 |
|  | PD-L2 | OID00831 | Q9BQ51 | 28 | 0.0014 | 0.0126 |
|  | CD27 | OID00800 | P26842 | 31 | 0.0022 | 0.0180 |
|  | CXCL10 | OID00807 | P02778 | 32 | 0.0025 | 0.0188 |
|  | CXCL9 | OID00771 | Q07325 | 33 | 0.0029 | 0.0188 |
|  | CD83 | OID00841 | Q01151 | 33 | 0.0029 | 0.0188 |
|  | GZMA | OID00804 | P12544 | 34 | 0.0033 | 0.0201 |
|  | MIC-A/B | OID00820 | Q29983.Q29980 | 36 | 0.0043 | 0.0246 |
|  | TNFRSF9 | OID00753 | Q07011 | 40 | 0.0071 | 0.0344 |
|  | CD28 | OID00793 | P10747 | 40 | 0.0071 | 0.0344 |
|  | MMP12 | OID00829 | P39900 | 40 | 0.0071 | 0.0344 |
|  | PDCD1 | OID00791 | Q15116 | 41 | 0.0080 | 0.0368 |
|  | IL10 | OID00809 | P22301 | 42 | 0.0090 | 0.0395 |
|  | NOS3 | OID00777 | P29474 | 44 | 0.0113 | 0.0454 |
|  | IL12RB1 | OID00835 | P42701 | 44 | 0.0113 | 0.0454 |
|  | TIE2 | OID00754 | Q02763 | 45 | 0.0127 | 0.0467 |
|  | TWEAK | OID00789 | O43508 | 45 | 0.0127 | 0.0467 |
|  | ADGRG1 | OID00764 | Q9Y653 | 48 | 0.0175 | 0.0598 |
|  | CD8A | OID00772 | P01732 | 48 | 0.0175 | 0.0598 |
|  | IL6 | OID00763 | P05231 | 50 | 0.0216 | 0.0709 |
|  | DCN | OID00817 | P07585 | 53 | 0.0290 | 0.0918 |
|  | VEGFR-2 | OID00780 | P35968 | 54 | 0.0319 | 0.0918 |
|  | CCL19 | OID00794 | Q99731 | 54 | 0.0319 | 0.0918 |
|  | TNFRSF4 | OID00819 | P43489 | 54 | 0.0319 | 0.0918 |
|  | KLRD1 | OID00839 | Q13241 | 56 | 0.0384 | 0.1072 |
|  | CD70 | OID00808 | P32970 | 59 | 0.0502 | 0.1358 |
|  | MCP-1 | OID00765 | P13500 | 61 | 0.0595 | 0.1521 |
|  | CSF-1 | OID00843 | P09603 | 61 | 0.0595 | 0.1521 |
|  | IFN-gamma | OID05552 | P01579 | 62 | 0.0646 | 0.1608 |
|  | CCL17 | OID00821 | Q92583 | 64 | 0.0760 | 0.1792 |
|  | ICOSLG | OID00828 | O75144 | 64 | 0.0760 | 0.1792 |
|  | ANGPT2 | OID00822 | O15123 | 65 | 0.0822 | 0.1890 |
|  | CAIX | OID00773 | Q16790 | 66 | 0.0888 | 0.1945 |
|  | TNFRSF12A | OID00810 | Q9NP84 | 66 | 0.0888 | 0.1945 |
|  | ANGPT1 | OID00760 | Q15389 | 67 | 0.0958 | 0.2050 |
|  | CXCL13 | OID00830 | O43927 | 68 | 0.1032 | 0.2158 |
|  | CD40-L | OID00756 | P29965 | 71 | 0.1281 | 0.2618 |
|  | PDGF subunit B | OID00790 | P01127 | 72 | 0.1373 | 0.2687 |
|  | PD-L1 | OID00799 | Q9NZQ7 | 72 | 0.1373 | 0.2687 |
|  | MCP-4 | OID00768 | Q99616 | 73 | 0.1470 | 0.2817 |
|  | CXCL11 | OID00767 | O14625 | 75 | 0.1678 | 0.3151 |
|  | FASLG | OID00792 | P48023 | 76 | 0.1790 | 0.3293 |
|  | EGF | OID00759 | P01133 | 79 | 0.2157 | 0.3816 |
|  | MCP-2 | OID00795 | P80075 | 79 | 0.2157 | 0.3816 |
|  | IL7 | OID00761 | P13232 | 80 | 0.2290 | 0.3975 |
|  | Gal-9 | OID00779 | O00182 | 81 | 0.2428 | 0.4062 |
|  | CXCL12 | OID00824 | P48061 | 81 | 0.2428 | 0.4062 |
|  | CXCL5 | OID00801 | P42830 | 85 | 0.3038 | 0.4904 |
|  | CCL23 | OID00811 | P55773 | 85 | 0.3038 | 0.4904 |
|  | TNFRSF21 | OID00818 | O75509 | 86 | 0.3205 | 0.5083 |
|  | IL15 | OID05551 | P40933 | 87 | 0.3377 | 0.5177 |
|  | CASP-8 | OID00827 | Q14790 | 87 | 0.3377 | 0.5177 |
|  | PTN | OID00823 | P21246 | 90 | 0.3926 | 0.5921 |
|  | ARG1 | OID00815 | P05089 | 40 | 0.4631 | 0.6794 |
|  | LAP TGF-beta-1 | OID00785 | P01137 | 94 | 0.4733 | 0.6794 |
|  | CD244 | OID00758 | Q9BZW8 | 95 | 0.4948 | 0.6794 |
|  | FGF2 | OID00770 | P09038 | 95 | 0.4948 | 0.6794 |
|  | MMP7 | OID00814 | P09237 | 95 | 0.4948 | 0.6794 |
|  | CCL20 | OID00837 | P78556 | 95 | 0.4948 | 0.6794 |
|  | CD4 | OID00776 | P01730 | 96 | 0.5168 | 0.6991 |
|  | Gal-1 | OID00798 | P09382 | 97 | 0.5392 | 0.7086 |
|  | IL13 | OID00836 | P35225 | 97 | 0.5392 | 0.7086 |
|  | CXCL1 | OID00786 | P09341 | 98 | 0.5621 | 0.7283 |
|  | IL-1 alpha | OID00757 | P01583 | 43 | 0.5830 | 0.7450 |
|  | MUC-16 | OID05549 | Q8WXI7 | 100 | 0.6091 | 0.7573 |
|  | IL2 | OID00778 | P60568 | 100 | 0.6091 | 0.7573 |
|  | CCL3 | OID00813 | P10147 | 101 | 0.6333 | 0.7768 |
|  | IL5 | OID00802 | P05113 | 102 | 0.6578 | 0.7860 |
|  | CX3CL1 | OID00806 | P78423 | 102 | 0.6578 | 0.7860 |
|  | KIR3DL1 | OID05550 | P43629 | 103 | 0.6827 | 0.8053 |
|  | TNFSF14 | OID00787 | O43557 | 104 | 0.7079 | 0.8141 |
|  | HO-1 | OID00805 | P09601 | 104 | 0.7079 | 0.8141 |
|  | IL33 | OID00788 | O95760 | 106 | 0.7593 | 0.8416 |
|  | LAMP3 | OID00826 | Q9UQV4 | 106 | 0.7593 | 0.8416 |
|  | VEGFA | OID00832 | P15692 | 106 | 0.7593 | 0.8416 |
|  | HGF | OID00803 | P14210 | 107 | 0.7854 | 0.8602 |
|  | IL8 | OID00752 | P10145 | 109 | 0.8382 | 0.9072 |
|  | ADA | OID00775 | P00813 | 110 | 0.8649 | 0.9252 |
|  | CCL4 | OID00796 | P13236 | 111 | 0.8917 | 0.9394 |
|  | IL4 | OID00833 | P05112 | 82 | 0.8986 | 0.9394 |
|  | CD40 | OID00781 | P25942 | 113 | 0.9457 | 0.9668 |
|  | IL18 | OID00782 | Q14116 | 113 | 0.9457 | 0.9668 |
|  | MCP-3 | OID00755 | P80098 | 115 | 1.0000 | 1.0000 |
|  | PGF | OID00762 | P49763 | 115 | 1.0000 | 1.0000 |
| OT_96IR | LAG3 | OID01023 | P18627 | 19 | 0.0003 | 0.0269 |
|  | CLEC6A | OID01017 | Q6EIG7 | 33 | 0.0029 | 0.1314 |
|  | SIT1 | OID01013 | Q9Y3P8 | 40 | 0.0071 | 0.2178 |
|  | LILRB4 | OID00965 | Q8NHJ6 | 47 | 0.0158 | 0.2821 |
|  | FCRL6 | OID01006 | Q6DN72 | 49 | 0.0195 | 0.2821 |
|  | SH2D1A | OID01002 | O60880 | 50 | 0.0216 | 0.2821 |
|  | IL10 | OID00993 | P22301 | 53 | 0.0290 | 0.2821 |
|  | IL5 | OID01024 | P05113 | 53 | 0.0290 | 0.2821 |
|  | DPP10 | OID00956 | Q8N608 | 54 | 0.0319 | 0.2821 |
|  | NCR1 | OID01007 | O76036 | 54 | 0.0319 | 0.2821 |
|  | CD28 | OID00977 | P10747 | 56 | 0.0384 | 0.2821 |
|  | IL12RB1 | OID01019 | P42701 | 56 | 0.0384 | 0.2821 |
|  | MASP1 | OID01014 | P48740 | 57 | 0.0421 | 0.2821 |
|  | KLRD1 | OID00995 | Q13241 | 58 | 0.0460 | 0.2821 |
|  | IFNLR1 | OID01010 | Q8IU57 | 58 | 0.0460 | 0.2821 |
|  | GLB1 | OID00937 | P16278 | 62 | 0.0646 | 0.3499 |
|  | EDAR | OID00946 | Q9UNE0 | 62 | 0.0646 | 0.3499 |
|  | CD83 | OID01025 | Q01151 | 63 | 0.0701 | 0.3585 |
|  | IL6 | OID00947 | P05231 | 69 | 0.1111 | 0.5109 |
|  | ITM2A | OID00968 | O43736 | 69 | 0.1111 | 0.5109 |
|  | CLEC4A | OID00951 | Q9UMR7 | 71 | 0.1281 | 0.5512 |
|  | CLEC4C | OID00949 | Q8WTT0 | 72 | 0.1373 | 0.5512 |
|  | PTH1R | OID00978 | Q03431 | 74 | 0.1571 | 0.5512 |
|  | FCRL3 | OID00984 | Q96P31 | 74 | 0.1571 | 0.5512 |
|  | MGMT | OID00990 | P16455 | 74 | 0.1571 | 0.5512 |
|  | TRIM5 | OID00957 | Q9C035 | 75 | 0.1678 | 0.5512 |
|  | ITGB6 | OID01026 | P18564 | 75 | 0.1678 | 0.5512 |
|  | FAM3B | OID01001 | P58499 | 76 | 0.1790 | 0.5512 |
|  | KPNA1 | OID01022 | P52294 | 76 | 0.1790 | 0.5512 |
|  | PIK3AP1 | OID00997 | Q6ZUJ8 | 77 | 0.1907 | 0.5512 |
|  | IRAK4 | OID00940 | Q9NWZ3 | 78 | 0.2029 | 0.5512 |
|  | MILR1 | OID00971 | Q7Z6M3 | 78 | 0.2029 | 0.5512 |
|  | CLEC7A | OID01016 | Q9BXN2 | 78 | 0.2029 | 0.5512 |
|  | HCLS1 | OID00942 | P14317 | 79 | 0.2157 | 0.5512 |
|  | HNMT | OID00969 | P50135 | 79 | 0.2157 | 0.5512 |
|  | TANK | OID01020 | Q92844 | 79 | 0.2157 | 0.5512 |
|  | PRDX5 | OID00955 | P30044 | 80 | 0.2290 | 0.5694 |
|  | DAPP1 | OID01011 | Q9UN19 | 81 | 0.2428 | 0.5879 |
|  | FXYD5 | OID00962 | Q96DB9 | 82 | 0.2572 | 0.6068 |
|  | IRAK1 | OID00950 | P51617 | 83 | 0.2722 | 0.6156 |
|  | GALNT3 | OID00961 | Q14435 | 84 | 0.2877 | 0.6156 |
|  | SPRY2 | OID00998 | O43597 | 84 | 0.2877 | 0.6156 |
|  | ARNT | OID01000 | P27540 | 84 | 0.2877 | 0.6156 |
|  | CKAP4 | OID00985 | Q07065 | 85 | 0.3038 | 0.6353 |
|  | TPSAB1 | OID00941 | Q15661 | 86 | 0.3205 | 0.6552 |
|  | PSIP1 | OID00938 | O75475 | 88 | 0.3554 | 0.6957 |
|  | NF2 | OID00981 | P35240 | 88 | 0.3554 | 0.6957 |
|  | FGF2 | OID00954 | P09038 | 90 | 0.3926 | 0.7258 |
|  | AREG | OID01009 | P15514 | 90 | 0.3926 | 0.7258 |
|  | DGKZ | OID00948 | Q13574 | 91 | 0.4120 | 0.7258 |
|  | ICA1 | OID01003 | Q05084 | 91 | 0.4120 | 0.7258 |
|  | TRAF2 | OID00963 | Q12933 | 92 | 0.4319 | 0.7258 |
|  | CNTNAP2 | OID00943 | Q9UHC6 | 93 | 0.4524 | 0.7258 |
|  | KRT19 | OID00967 | P08727 | 93 | 0.4524 | 0.7258 |
|  | EIF5A | OID00975 | P63241 | 93 | 0.4524 | 0.7258 |
|  | BACH1 | OID00996 | O14867 | 93 | 0.4524 | 0.7258 |
|  | DCTN1 | OID00958 | Q14203 | 94 | 0.4733 | 0.7258 |
|  | EIF4G1 | OID00976 | Q04637 | 94 | 0.4733 | 0.7258 |
|  | SH2B3 | OID00983 | Q9UQQ2 | 94 | 0.4733 | 0.7258 |
|  | PRKCQ | OID00989 | Q04759 | 94 | 0.4733 | 0.7258 |
|  | CXCL12 | OID01008 | P48061 | 95 | 0.4948 | 0.7463 |
|  | NTF4 | OID00966 | P34130 | 96 | 0.5168 | 0.7546 |
|  | HEXIM1 | OID00987 | O94992 | 96 | 0.5168 | 0.7546 |
|  | DFFA | OID01004 | O00273 | 97 | 0.5392 | 0.7751 |
|  | DDX58 | OID01018 | O95786 | 98 | 0.5621 | 0.7956 |
|  | DCBLD2 | OID01005 | Q96PD2 | 99 | 0.5854 | 0.8122 |
|  | ZBTB16 | OID00939 | Q05516 | 100 | 0.6091 | 0.8122 |
|  | CCL11 | OID00970 | P51671 | 100 | 0.6091 | 0.8122 |
|  | PLXNA4 | OID00982 | Q9HCM2 | 100 | 0.6091 | 0.8122 |
|  | HSD11B1 | OID00980 | P28845 | 102 | 0.6578 | 0.8524 |
|  | BTN3A2 | OID01027 | P78410 | 102 | 0.6578 | 0.8524 |
|  | BIRC2 | OID00979 | Q13490 | 103 | 0.6827 | 0.8604 |
|  | STC1 | OID00999 | P52823 | 103 | 0.6827 | 0.8604 |
|  | CLEC4D | OID00988 | Q8WXI8 | 105 | 0.7335 | 0.9119 |
|  | TRIM21 | OID00964 | P19474 | 106 | 0.7593 | 0.9191 |
|  | NFATC3 | OID00973 | Q12968 | 106 | 0.7593 | 0.9191 |
|  | IRF9 | OID00945 | Q00978 | 108 | 0.8117 | 0.9698 |
|  | PPP1R9B | OID00936 | Q96SB3 | 109 | 0.8382 | 0.9761 |
|  | CDSN | OID00960 | Q15517 | 109 | 0.8382 | 0.9761 |
|  | TREM1 | OID00991 | Q9NP99 | 110 | 0.8649 | 0.9884 |
|  | CLEC4G | OID00944 | Q6UXB4 | 111 | 0.8917 | 0.9884 |
|  | SRPK2 | OID00994 | P78362 | 111 | 0.8917 | 0.9884 |
|  | LAMP3 | OID01015 | Q9UQV4 | 111 | 0.8917 | 0.9884 |
|  | PRDX1 | OID00952 | Q06830 | 112 | 0.9187 | 0.9944 |
|  | PRDX3 | OID00953 | P30048 | 112 | 0.9187 | 0.9944 |
|  | EGLN1 | OID00972 | Q9GZT9 | 113 | 0.9457 | 0.9945 |
|  | PADI2 | OID01012 | Q9Y2J8 | 113 | 0.9457 | 0.9945 |
|  | LY75 | OID00974 | O60449 | 114 | 0.9729 | 0.9945 |
|  | JUN | OID00986 | P05412 | 114 | 0.9729 | 0.9945 |
|  | CXADR | OID00992 | P78310 | 114 | 0.9729 | 0.9945 |
|  | ITGA6 | OID00959 | P23229 | 115 | 1.0000 | 1.0000 |
|  | ITGA11 | OID01021 | Q9UKX5 | 115 | 1.0000 | 1.0000 |
